# Spatial transcriptomic profiling uncovers the molecular effects of the neurotoxicant polychlorinated biphenyls (PCBs) in the brains of adult mice

**DOI:** 10.1101/2025.07.24.665575

**Authors:** Budhaditya Basu, Nicole M. Breese, Sal Lombardi, Hui Wang, Xueshu Li, Destiny Tiburcio, Zachary Niemasz, Stacy E. Beyer, Laura E. Dean, Rachel F. Marek, Michal Toborek, Hans-Joachim Lehmler, Snehajyoti Chatterjee

## Abstract

Environmental toxicants, such as polychlorinated biphenyls (PCBs), are highly stable synthetic organic compounds that are present in air, water, and soil. PCBs have been identified in post-mortem human brains of individuals with neurodegenerative diseases, indicating a possible link between environmental factors and disease risk. Research has revealed an association between PCB exposure and cognitive decline. Therefore, it is crucial to evaluate how PCB mixtures relevant to humans affect brain function and cognition. To investigate the effects of PCBs on memory and transcriptomic profiles, we exposed adult male C57BL/6J mice orally to a synthetic PCB mixture daily. After seven weeks of exposure, the mice were assessed in a spatial object recognition task (SOR) to evaluate long-term spatial memory. Our findings showed that mice exposed to PCBs exhibited deficits in long-term spatial memory. To examine the molecular effects of PCB on the brain, we used a spatial transcriptomics technique to analyze gene expression changes in five brain regions: the hippocampus, neocortex, thalamus, caudal putamen, and fiber tracts. Our analysis of spatial gene expression revealed the molecular signatures influenced by PCB in these susceptible brain regions of mice. Network analysis suggests that these changes are associated with higher chlorinated PCBs present in the brain. Additionally, we show that PCB exposure disrupts the expression of tight junction proteins, which are crucial for maintaining the integrity of the blood-brain barrier (BBB). Thus, our results offer mechanistic insights into how PCB exposure affects brain function and cognition.

## Introduction

Epidemiological studies suggest environmental contaminants, especially neurotoxicants, are associated with Alzheimer’s disease and related dementias (ADRD) and other neurodegenerative diseases [1, 2]. Polychlorinated biphenyls (PCBs) are a group of synthetic organic compounds and are associated with neurotoxicity [3, 4]. PCB manufacturing began in 1929, and due to their properties, such as chemical stability, non-flammability, and electrical insulating properties, PCBs were widely used in various applications, including dielectric fluids, capacitors, transformers, pesticides, lubricants, and construction materials. In the United States, PCB manufacturing was banned in 1979 due to environmental and health concerns. However, PCBs continue to be produced inadvertently as byproducts of the manufacturing of consumer products, such as paints and silicone rubber [5, 6]. They are released into the environment and remain an ongoing public health threat due to their environmental persistence and the fact that they bioaccumulate and biomagnify in the food chain. PCBs are lipophilic compounds stored in milk fat and adipose tissue throughout the body [7, 8]. PCBs have been found in post-mortem human brains [9], and elevated levels of PCBs have been associated with reduced cognitive performance in older adults [10], suggesting a link between PCB-mediated neurotoxicity and memory performance in dementia. However, these factors are challenging to study in human populations because of the long time between exposure and the onset of ADRD. Therefore, the PCB-mediated impact on the molecular signature in the brain and behavior remains unclear.

The hippocampus and neocortex are critical brain regions closely associated with spatial memory consolidation [11, 12], and gene expression in the hippocampus [13, 14] and cortex [15–17] play an important role in learning and memory. Using a spatial transcriptomics approach and deep learning computational tool, we recently showed that learning in a spatial memory task increases predicted neuronal activity and expression of memory-related genes in the hippocampus and neocortical regions [14, 18]. The hippocampus and neocortex are also brain regions vulnerable to various neuropathologies, including ADRD [19, 20] and PCB induced neurotoxicity [9, 21, 22]. Thus, understanding the gene expression signature in the hippocampus and neocortex following PCB-induced neurotoxicity could provide mechanistic insights into how neurotoxicants, such as PCB, impact brain function. However, such an in-depth transcriptomic analysis to study the impact of PCB on different brain regions has not been performed to date.

In this study, we investigated the impact of a synthetic human-relevant PCB mixture (HR-PCB) on spatial memory and brain transcriptomics in adult male C57BL/6J mice. This synthetic PCB mixture approximates the average PCB mass profile found in postmortem human brain tissues. We found that exposure to this PCB mixture impaired long-term spatial memory. Next, we applied a state-of-the-art spatial transcriptomics approach to examine gene expression changes in response to PCB exposure in five brain regions: hippocampus, neocortex, thalamus, fiber tracts, and caudoputamen. Furthermore, we demonstrate that PCB exposure affects the expression of key genes related to memory and synaptic plasticity. Network analysis revealed that mostly higher chlorinated PCB congeners were associated with specific differentially expressed genes identified in the transcriptomics analyses. We also showed that PCB exposure affects the expression and levels of proteins involved in BBB integrity and permeability. Collectively, we demonstrate the molecular signature of PCB-mediated neurotoxicity in adult mice.

## Results

### PCB impairs long-term spatial memory in adult mice

We exposed mice to a human-relevant PCB (HR-PCB) mixture designed to simulate the average profile found in post-mortem human brain samples (**Supplemental Figure S1**). Studies in mouse models of ADRD show spatial memory impairment in spatial object recognition (SOR) tasks [13], a behavioral test frequently used to assess hippocampus-dependent spatial memory. Given the link between spatial memory loss and ADRD, we assessed the impact of the HR-PCB mixture (**Supplemental Fig. S1**) on spatial memory consolidation in adult mice. Briefly, male mice received daily oral doses of PCB mix (6 mg/kg/d) or vehicle alone for seven weeks. This dose of HR-PCB had no impact on the overall body and brain weight of these animals (**Supplemental Fig. S2**). After completing HR-PCB dosing, mice were habituated in an open field and trained in a SOR task (**Fig. 1A**). During the habituation session, both HR-PCB and vehicle-exposed mice showed similar time spent in the inner and outer zones of the open field arena (**Fig. 1B**). This indicates that the administered dose of HR-PCB does not affect any anxiety-like behavior in adult mice. For the spatial memory assessment, mice were trained in the SOR task, where animals were exposed to the open field with two identical objects in specific spatial locations. The animals were trained to learn the spatial location of the objects, and their long-term memory was tested 24 hours later. During the test session, one object was relocated to a new spatial location to assess long-term spatial memory (**Fig. 1C**). We found that vehicle-exposed mice showed discrimination towards the moved object during the test session compared to the training session, suggesting normal long-term spatial memory (**Fig. 1C**). Importantly, HR-PCB-exposed mice could not discriminate between the objects during the test session (**Fig. 1C**), despite no differences in total exploration time (**Fig. 1D**). These results show that exposure to PCBs negatively impacted the long-term spatial memory of adult male mice.

**Figure 1.**
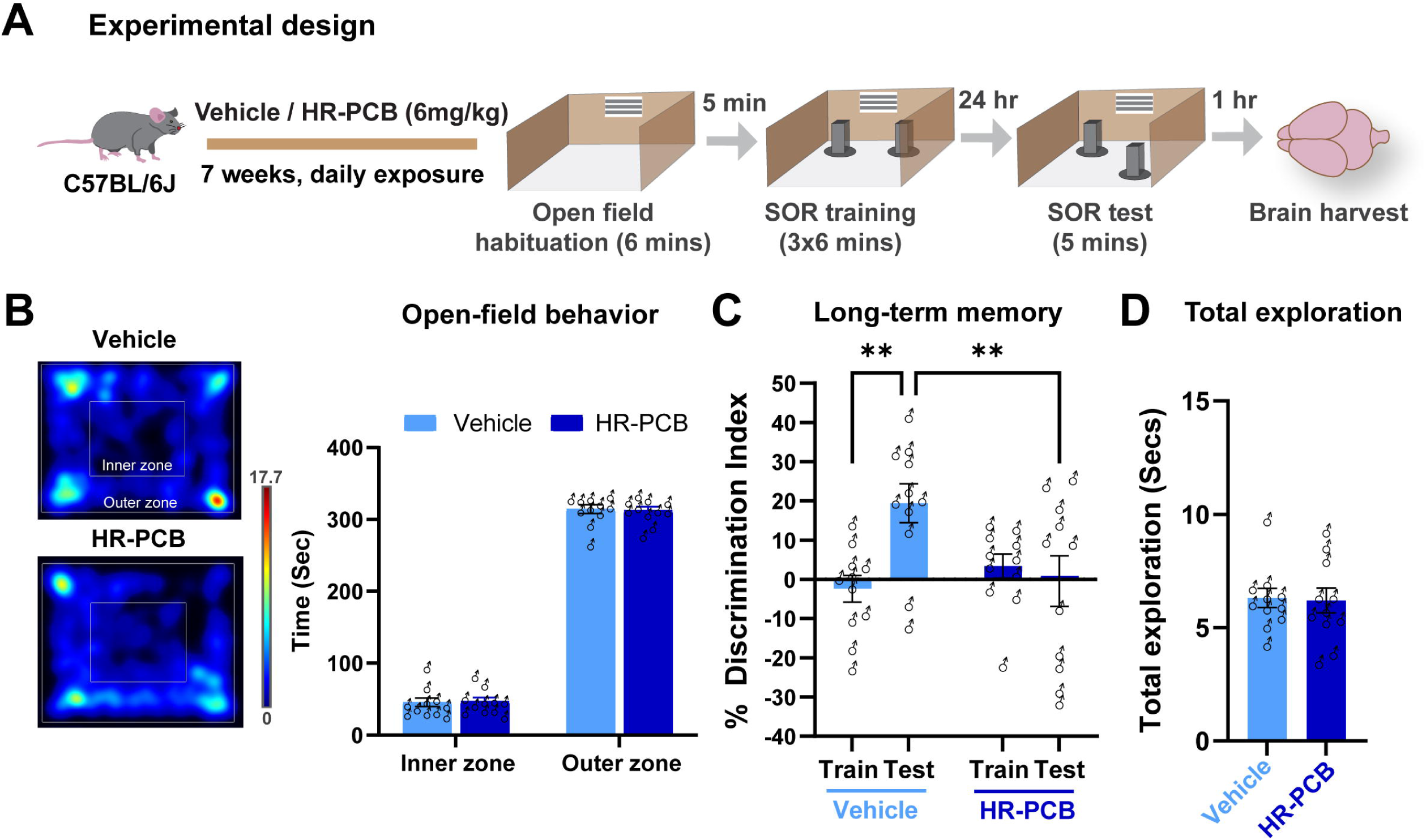
Long-term spatial memory assessment of wild-type mice exposed to the HR-PCB mixture. **A.** Experimental design. HR-PCB or vehicle was orally administered to male C57BL/6J mice for 7 weeks. After completion of the dosing, mice were trained in the spatial object recognition (SOR) task and tested for long-term memory 24 hours later. **B.** Time spent in the inner and outer zone of the open field during the habituation session. **C.** Long-term spatial memory assessment of mice by evaluating discrimination for the displaced object (DO) in the SOR task. 2-way Anova: Significant session (Train-Test) x exposure (PCB-vehicle) interaction: F_(1, 20)_= 8.697, *p*=0.0079. Sidak’s multiple comparison tests: vehicle (train) vs. vehicle (test): \*\**p*=0.0041, and vehicle (test) vs. PCB (test): \*\**p*=0.0092. Error bars represent ± SEM. Vehicle (n=11), and HR-PCB (n=11). **D.** Total exploration of the objects during the test session.

### PCB exposure leads to high-chlorinated PCB congeners enrichment in mouse brains

PCBs have been detected in post-mortem human brain tissues [9, 23] and tissues from PCB-exposed laboratory animals [24]. In mice exposed to a complex PCB mixture, specific PCB congeners present in the brain have been directly linked to changes in the brain transcriptome [24]. These changes may be ultimately associated with behavioral outcomes. We characterized the PCB profiles and levels in the brains of mice exposed to the HR-PCB mixture (n = 4 per group) using a liquid-liquid extraction method, followed by gas chromatography-tandem mass spectrometry (GC-MS/MS) analysis. This congener-specific method can identify and quantify all 209 PCB congeners as 173 peaks of individual or co-eluting PCB congeners. Overall, the PCB profiles in the mouse brain differed from the original exposed HR-PCB mixture profile (similarity coefficient cos θ = 0.54), with enrichment in the high-chlorinated PCB congeners [25](**Fig. 2**), possibly due to the more rapid metabolism of lower-chlorinated PCB congeners. A total of 69 individual or co-eluting PCB congeners were detected, with a detection frequency of over 50%. PCB congener levels ranged from non-detectable to 500 ng/g ww. The total ∑PCB burden in the brain was 3360 ng/g ww, dominated by PCB153+157 (500 ± 100 ng/g), followed by PCB129+138+163 (490 ± 90 ng/g) and PCB180+193 (450 ± 100 ng/g). Examining the homolog levels, the majority of observed PCB congeners ranged from penta- to nonachlorobiphenyls. Surprisingly, neither mono nor dichlorobiphenyls were detected in the brain, while PCB20+28 was the only trichlorobiphenyl detected. Moreover, to estimate the levels of dioxin-like PCBs, we calculated the toxic equivalency (WHO-TEQ), used for human risk assessment, as 0.013 ng/g ww using the revised toxic equivalency factors (TEF) [26], with major contributions from PCB118 and PCB105.

**Figure 2.**
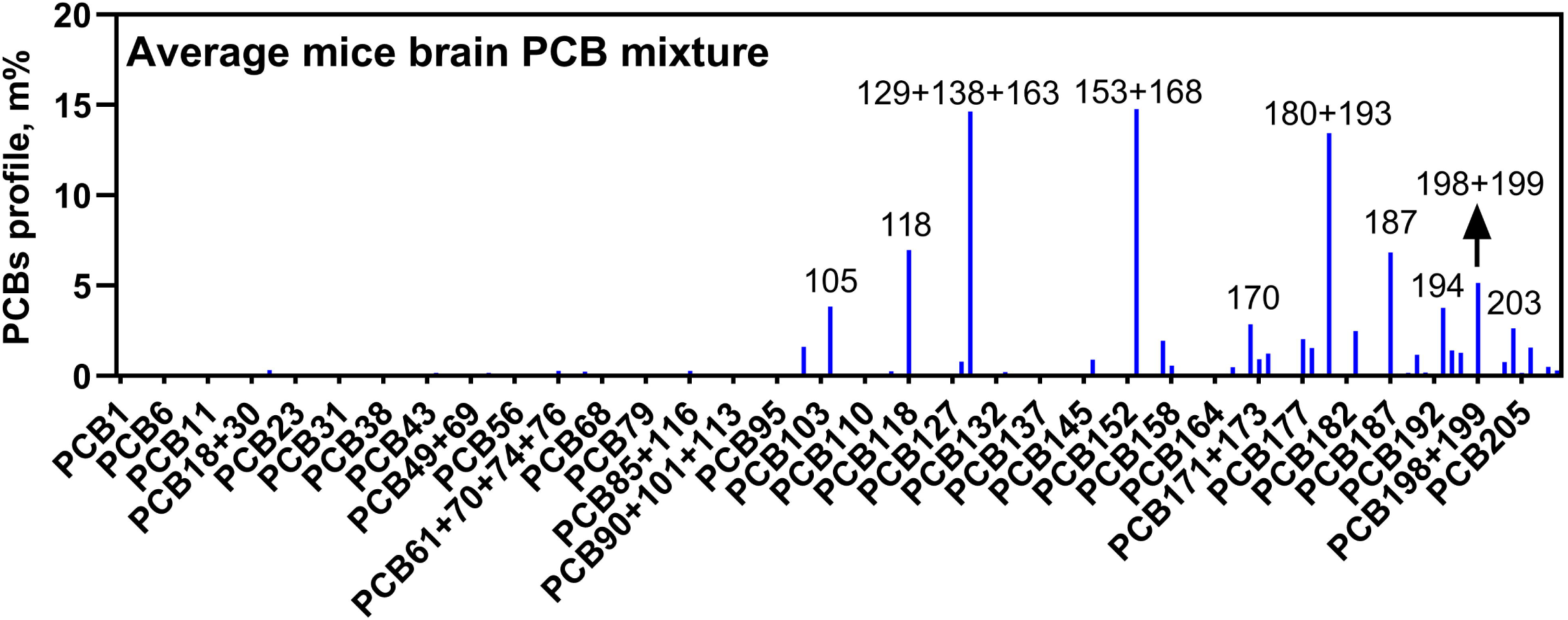
PCB profile in mouse brain. Average PCB profile in mouse brains from this study (n = 4 animals per group). The PCB profiles were determined by GC-MS/MS.

### PCB mediates distinct gene expression signatures in different brain regions

Spatial transcriptomics is a powerful tool for determining gene expression changes at a high spatial resolution. Using spatial transcriptomics approaches, we have previously shown gene expression changes across brain sub-regions during spatial memory consolidation [14, 18]. Therefore, we investigated spatial gene expression changes impacted by the HR-PCB mixture. Brain sections from PCB- and vehicle-exposed male mice, collected one hour after the memory test session, were processed for spatial transcriptomic analyses using the Visium platform. Based on spatial location and marker gene expression, our analysis identified five major brain sub-regions (neocortex, hippocampus, thalamus, fiber tracts, and caudoputamen) (**Fig. 3A-D**). Importantly, our findings indicate that HR-PCB mixture and vehicle-exposed mice have a similar distribution of barcode spots in the hippocampus and fiber tracts (**Fig. 3E**). However, there are a reduced proportion of barcode spots in the neocortex and caudoputamen, and an increased proportion of barcode spots in the thalamus region of the PCB-treated group. (**Fig. 3E**). A Fisher’s exact test demonstrated notable variations in the proportion of barcoded spots within the neocortex, caudoputamen, and thalamus. In contrast, no significant difference was observed in the hippocampal and fiber tract regions between the PCB and vehicle exposed groups. These differences are likely attributable to the method of preparation and placement of the coronal brain sections within the Visium capture area, which was centered on the hippocampal region (**Supplemental Fig. S3**).

**Figure 3.**
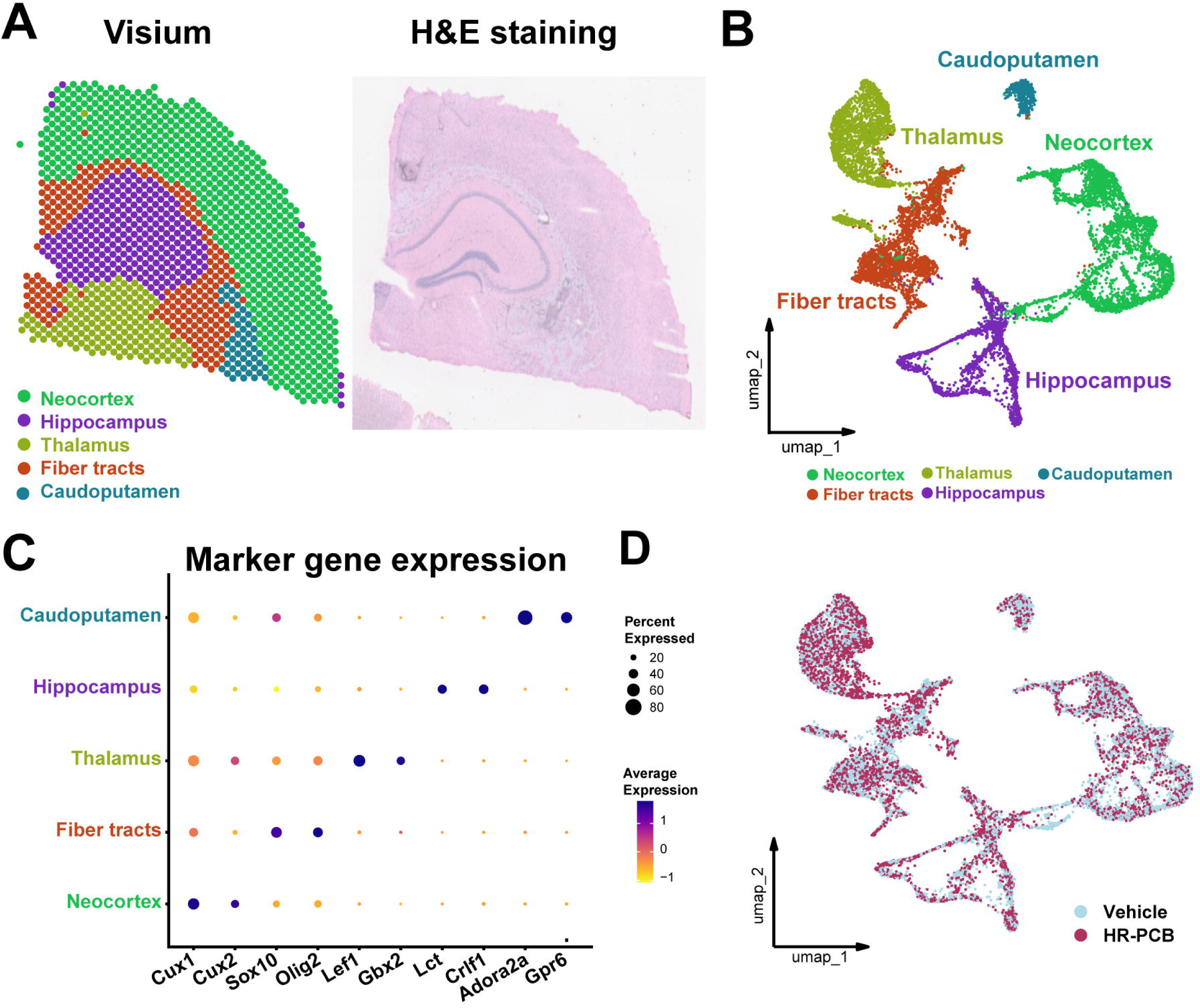
Spatial transcriptomics analysis of brain samples from wild-type mice exposed to HR-PCB mixture. **A.** Graph-based cluster identification was done at the spot level of samples. Spot color was assigned based on the transcriptomic signature using the Louvain clustering algorithm. Corresponding H&E histology staining of the Visium slice is also shown. **B.** UMAP plot based on the transcriptional signature of each spot. **C.** Dot plot showing expression levels of marker genes in annotated brain regions. The color of the dot plot shows the average expression of genes and dot size represents the percentage of spots expressing the gene of a given group. **D.** UMAP showing the clustering of barcoded spots from the vehicle and HR-PCB groups.

Differential gene expression analysis between HR-PCB-exposed and vehicle-exposed mice for each of the five brain regions found that the thalamus exhibited the greatest number of significant differentially expressed genes (DEGs) impacted by the HR-PCB mixture (177 DEGs), followed by the fiber tracts (162 DEGs), hippocampus (161 DEGs), neocortex (148 DEGs), and caudoputamen (72 DEGs) (**Fig. 4A**). Within these brain regions, PCB exposure resulted in 75 upregulated and 102 downregulated genes in the thalamus, 91 upregulated and 71 downregulated genes in fiber tracts, 116 upregulated and 45 downregulated genes in the hippocampus, 54 upregulated and 94 downregulated genes in the neocortex, and 43 upregulated and 29 downregulated genes in the caudoputamen (**Fig. 4A**).

**Figure 4.**
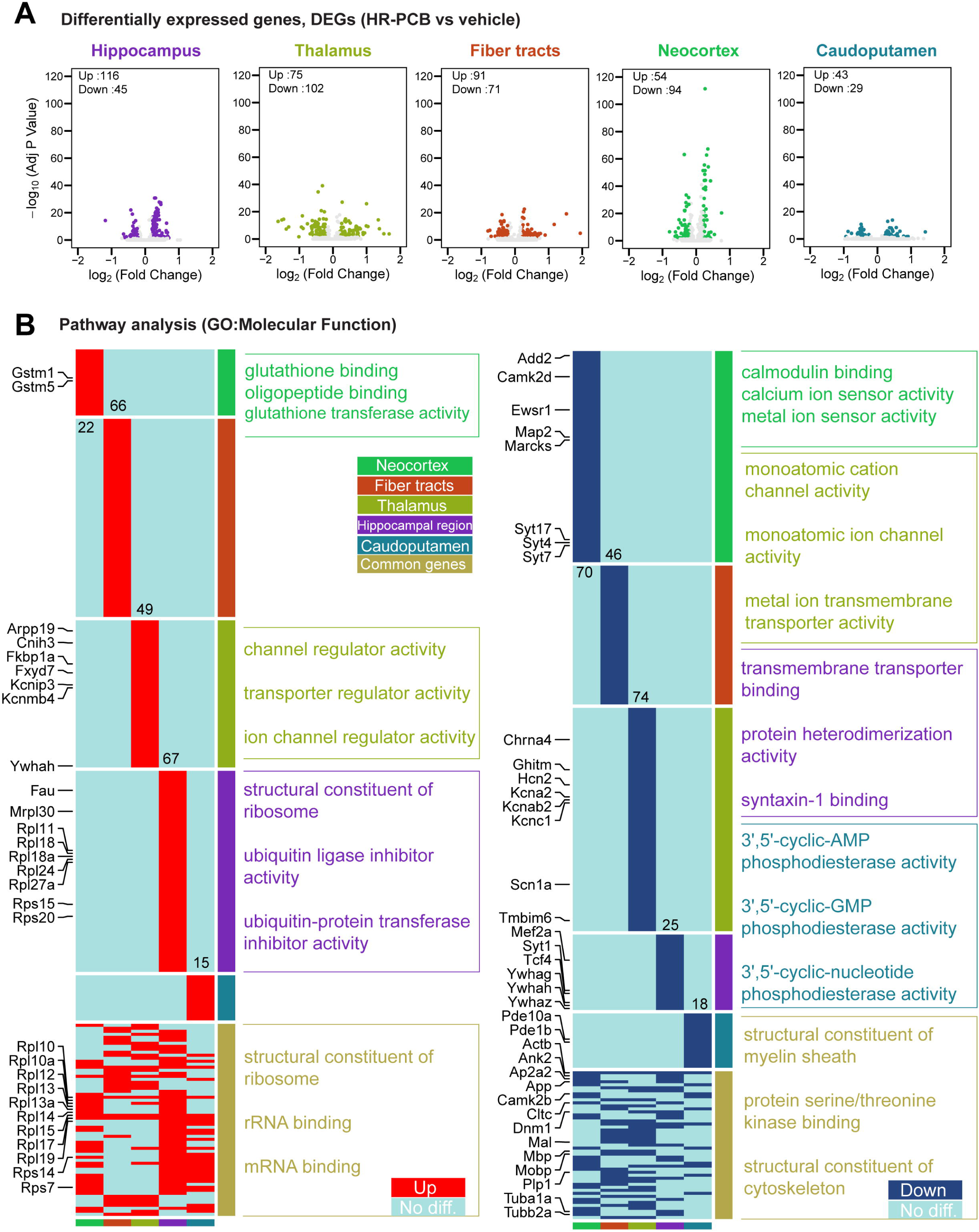
Genes impacted by HR-PCBs in the mouse brain. **A.** Volcano plots showing differentially expressed genes impacted by HR-PCB mixture in different brain regions: hippocampus, thalamus, fiber tracts, neocortex, and caudoputamen. **B.** Pathway analysis showing significantly enriched pathways (GO: Molecular function) (p.adjust < 0.01 & qvalue < 0.05) on unique or common genes upregulated or downregulated in five brain regions. The top three gene function annotations are shown. Upregulated and downregulated DEGs are shown in red and blue colors, respectively.

From the upregulated genes, we observed 64 shared genes upregulated in more than one subregion and 219 subregion-specific genes upregulated exclusively across the five brain subregions. Functional enrichment (Gene ontology: Molecular Function) analysis showed that 64 shared genes were related to structural constituent of the ribosome, rRNA binding and mRNA binding (*Rpl10, Rpl12, Rpl13*), indicating global changes in ribosomal biosynthesis machinery (**Fig. 4B**). For the 219 subregion-specific upregulated genes, different molecular functions were enriched in each subregion. For instance, glutathione binding and glutathione transferase activity related genes (*Gstm1, Gstm5*) [27] were upregulated in neocortex region potentially indicating upregulation of an antioxidative defense mechanism to combat exogenous PCB compounds. The thalamic region had DEGs associated with ion channel regulator activity, transporter regulator activity (*Arpp19, Cnih3, Fkbp1a, Fxyd7*). Specific DEGs related to the structural constituent of ribosome, ubiquitin ligase inhibitor activity, and ubiquitin-protein transferase inhibitor activity (*Rpl11, Rpl37, Rpl5, Rps15, Rps20*) were upregulated in the hippocampal region exhibiting a stress response.

For the downregulated DEGs, we observed 49 shared and 233 subregion-specific genes downregulated across five brain regions (**Fig. 4B**). The shared genes were related to the structural constituent of myelin sheath (*Mal, Mbp, Mobp, Plp1*), protein serine/threonine kinase binding ( *Ap2a2, App, Cltc, Dnm1*), and structural constituent of cytoskeleton (*Actb, Tubb2a, Ank2, Camk2b, Tuba1a*). We found that exposure to PCB impacted genes related to the calmodulin-binding pathway and calcium ion sensor activity in the neocortex (**Fig. 4B**). Notably, the genes within this pathway (*Camk2d*, *Map2*, *Ewsr1*, *Marcks*, *Syt7*, *Add2*) were downregulated in the PCB-exposed group. Some of the genes related to the calmodulin-binding pathway are linked to synaptic plasticity and long-term memory storage [28, 29]. The thalamus showed downregulation of genes in pathways associated with monoatomic cation channel activity and monoatomic ion channel activity (*Kcnc1, Kcnc3*). Additionally, DEGs associated with transmembrane transporter binding (*Ywhah, Ywhaz*) and protein heterodimerization activity (Mef2a, Syt1, Tcf4, Ywhah) were enriched in the hippocampal region. The caudoputamen was found to be enriched with 3’,5’-cyclic-AMP phosphodiesterase activity related genes (*Pde10a, Pde1b*). These results provide a broad spectrum of gene targets and pathways impacted by exposure to the HR-PCB mixture in the mouse brain.

Spatial transcriptomics enables the detection of gene expression changes across different brain regions, allowing for the investigation of distinct gene signatures within these regions. Therefore, we utilized an UpSet plot to compare the DEGs from each brain region to identify the distinct as well as overlapping gene expression signatures mediated by PCB exposure in these five brain regions. Given the roles of the hippocampus and neocortex in long-term memory consolidation and vulnerability to ADRD, we investigated the genes that were distinctly downregulated in only the hippocampus or the neocortex to provide mechanistic insights into PCB-mediated memory impairment in adult mice. We found that 25 genes were distinctly downregulated in the hippocampus and 70 genes in the neocortex (**Fig. 5A**). Among the 25 downregulated genes in the hippocampus, *Tcf4* has been previously shown to regulate synaptic plasticity and memory formation [30]. Next, we examined the genes downregulated by HR-PCB across different brain regions. We found 16 genes distinctly downregulated in the fiber tracts and thalamus (*Lars2*, *Cplx1*, *Gad1*, *Kif5a*, *Kif5b*, *Mal*, *Mbp*, *Cldn11*, *Plp1*, *Qk*, *Slc1a2*, *Lgi3*, *Trf*, *Ugt8a*, and *Rapgef4*), 10 genes in the neocortex and hippocampus (*Ank2*, *Ap2a2*, *Dpysl2*, *Napb*, *Spock1*, *Morf4l1*, *Tuba1a*, *Tubb2a*, *Cltc*, and *Dynll2*), 6 genes in caudoputamen and neocortex (*Arf3*, *CaMK2b*, *CaMKv*, *Mapk1*, *Ncdn*, and *Rgs7bp*), 4 genes in thalamus and hippocampus (*Atp1b1*, *Dnm1*, *Map1b*, and *Kcnc2*), 3 genes in neocortex and thalamus (*Syngr1*, *Rab3c*, and *Tspyl4*), 2 genes in neocortex and fiber tracts (*R3hdm4*, and *Pkp4*), 2 genes in caudoputamen and fiber tracts (*Ddn*, and *Ppp3ca*), and 2 genes in fiber tracts, thalamus and hippocampus (*Pvalb*, and *Atp1a3*) (**Fig. 5A**).

**Figure 5.**
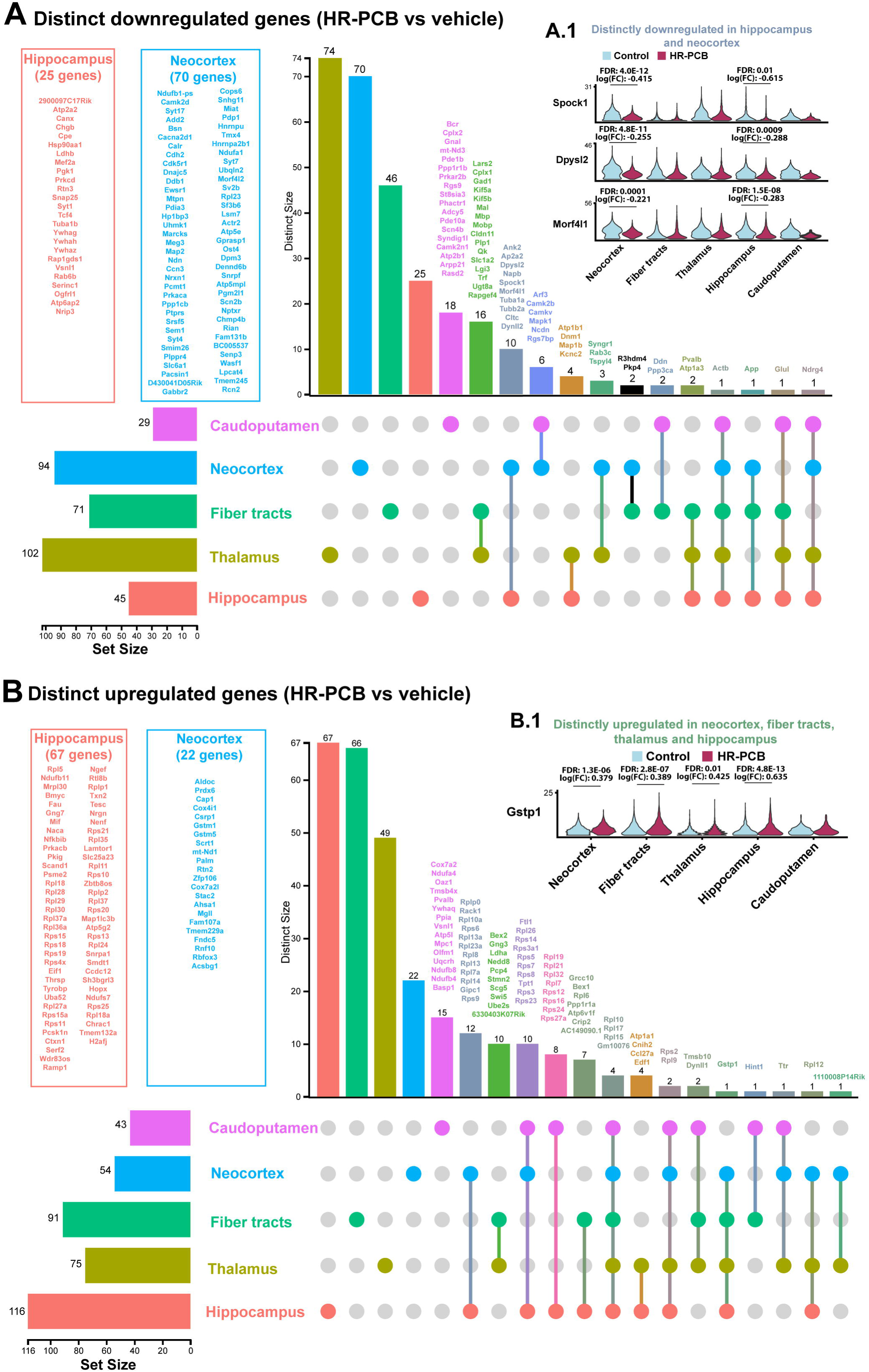
Distinct gene expression signature of HR-PCB exposure across five brain regions. **A.** Upset plot showing distinct and overlapping genes downregulated by PCB exposure in mouse brain sub-regions: Hippocampus, Thalamus, Neocortex, and Caudoputamen. ***A.1***. Violin plot showing the expression of *Spock1*, *Dpysl2*, and *Morf4l1* in mouse brains. **B.** Upset plot showing distinct and overlapping genes upregulated by PCB exposure in mouse brain sub-regions: Hippocampus, Thalamus, Neocortex, and Caudoputamen. ***B.1***. Violin plot showing the expression of *Gstp1* in the mouse brain.

*Dpysl2*, a gene downregulated distinctly in only the neocortex and hippocampus, is a synaptic-enriched protein [31] associated with dendritic spine formation and maturation [32], a process linked to memory consolidation. Importantly, brain-specific loss of Dpysl2 leads to hippocampus-dependent spatial memory impairment [33]. A recent study suggests that Dpysl2 also acts as an m^6^A reader, and the disruption of the interaction between Dpysl2 and Malat1 is linked to fear extinction memory loss [34]. Another gene downregulated in both the neocortex and hippocampus was *Morf4l1*. *Morf4l1* is part of the NuA4 histone acetyltransferase complex and Sin3b deacetylase complex. *Spock1*, also downregulated in the hippocampus and neocortex, is associated with blood-brain barrier (BBB) permeability. Previous reports suggest that PCB disrupts BBB integrity in mice [35] and rats [22], primarily by disrupting proteins related to tight junctions in the hippocampus, cerebrum, and cerebellum [22].

To study the genes upregulated following PCB exposure, we again used an UpSet plot to examine the genes across different brain regions (**Fig. 5B**). Sixty-seven genes were distinctly upregulated only in the hippocampus, and only 22 were upregulated in the neocortex. Comparing the distinctly upregulated genes across different brain regions, we found 12 genes distinctly upregulated in the neocortex and hippocampus (*Cox7a2*, *Ndufa4*, *Oaz1*, *Tmsb4x*, *Pvalb*, *Ywhaq*, *Ppia*, *Vsnl1*, *Atp5l*, *Mpc1*, *Olfm1*, *Uqcrh*, *Ndufb8*, *Ndufb4*, and *Basp1*), 10 genes in the fiber tracts and thalamus (*Bex2*, *Gng3*, *Nedd8*, *Pcp4*, *Stmn2*, *Scg5*, *Swi5*, *Ube2s*, and *6330403K07Rik*), 10 genes in the caudoputamen, neocortex, and hippocampus (*Ftl1*, *Rpl26*, *Rps14*, *Rps3a1*, *Rps5*, *Rps7*, *Rps8*, *Tpt1*, *Rps3,* and *Rps23*), 8 genes in caudoputamen and hippocampus (*Rpl19*, *Rpl21*, *Rpl32*, *Rpl7*, *Rps12*, *Rps16*, *Rps24*, and *Rps27a*), 7 genes in fiber tracts and hippocampus (*Grcc10*, *Bex1*, *Rpl6*, *Ppp1r1a*, *Atp6v1f*, *Crip2,* and *AC149090.1*), 4 genes in the caudoputamen, neocortex, fiber tracts, thalamus and hippocampus (*Rpl10*, *Rpl17*, *Rpl15* and *Gm10076*), 4 genes in thalamus and hippocampus (*Atp1a1*, *Cnih2*, *Ccl27a*, and *Edf1*), 2 genes in caudoputamen, neocortex, thalamus and hippocampus (*Rps2* and *Rpl9*), and 2 genes in caudoputamen, fiber tracts, and thalamus (*Tmsb10* and *Dynll1*).

Among the upregulated genes, Glutathione S-transferases (GSTs) are a group of proteins known to detoxify cells from endogenous and exogenous toxicants [36, 37]. Glutathione S-transferase P1 (Gstp1) was upregulated after PCB exposure in the neocortex, fiber tracts, thalamus, and hippocampus. Gstp1 is linked to detoxification and anti-oxidative damage [38], so, the upregulation of *Gstp1* in response to PCBs reflects the presence of toxicants in the brain tissue. Overall, our spatial transcriptomics analysis identified a unique gene expression signature mediated by PCB exposure in mouse brains.

### Interaction network analyses of the brain PCB spatial transcriptome revealed the association of specific genes with PCB congeners

We previously used network analyses with xMWAS to identify specific PCB congeners associated with differentially expressed genes identified by bulk RNA sequencing [39]. xMWAS provides integrative and association analysis between two datasets using partial least squares regression methods. Here, we utilized this platform to perform an interaction network analysis to identify associations between specific PCB congeners and specific DEGs identified in the spatial transcriptomic analysis [39]. We selected four genes affected by PCB exposure in the neocortex and hippocampus—*Dpysl2*, *Gstp1*, *Morf4l1*, and *Spock1*—and studied the relationships between these genes and individual PCBs in the mouse brain. Subnetworks centered on these genes were extracted from a comprehensive DEG network, which enabled the identification of PCB congeners linked to their altered expression.

In the neocortex, a total of 62 PCBs were identified to be closely connected to these four genes (*Morf4l1, Spock1, Dpysl2,* and *Gstp1*) (**Fig. 6A**). These PCBs were predominantly tetra to octachlorinated PCBs. All 62 PCB congeners identified in the PCB analysis were negatively correlated with *Morf4l1*. Twenty-four PCBs were negatively correlated with *Dpysl2*, thirteen with *Spock1*, and one PCB congener (PCB174) was positively correlated with *Gstp1*. Thirteen PCBs, mostly tetra to hexachlorinated congeners, were correlated with three genes (*Morf4l1, Spock1,* and *Dpysl2*). *Spock1* and *Dpysl2* were associated with tetra to hexachlorinated PCBs. Meanwhile, *Morf4l1* interacted with a wider range of PCBs, from tetra- to decachlorinated congeners.

**Figure 6.**
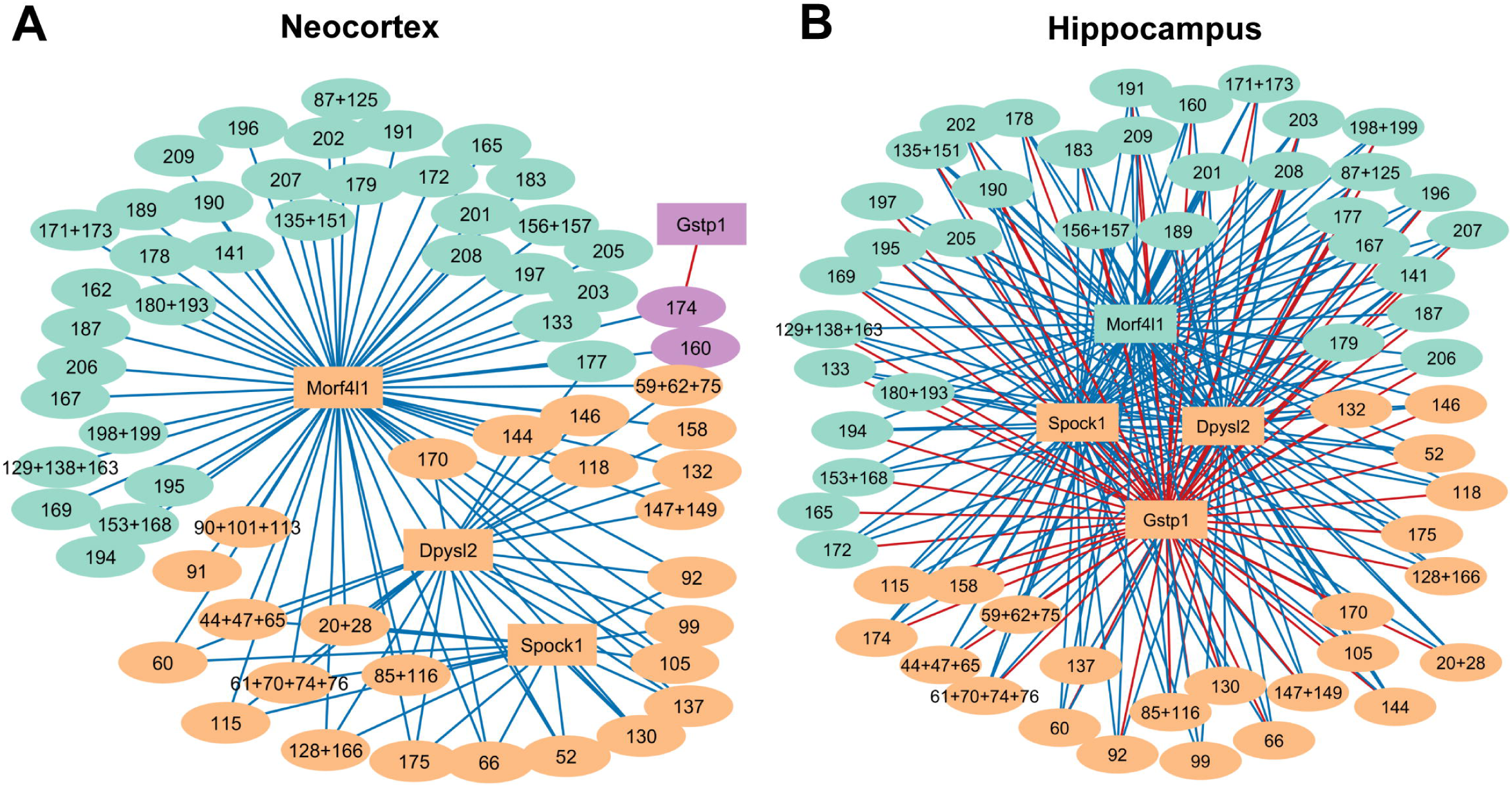
Interaction network analyses of genes and PCB congeners. xMWAS identified connections and clusters between brain HR-PCB levels and 4 selected DEGs identified in the (A) neocortex and (B) hippocampus. The analyses used a threshold of absolute correlation coefficients > 0.7 for neocortex, and > 0.9 for hippocampus. The p-value threshold for student’s t-test was < 0.05. Nodes in the same color indicate clusters. The node shape represents PCBs (ovals) and genes (rectangles). The line color indicates positive (red) and negative (blue) correlations.

In the hippocampus, a total of 59 polychlorinated biphenyls (PCBs) demonstrated strong associations with the four genes (*Morf4l1, Spock1, Dpysl2,* and *Gstp1*) (**Fig. 6B**). These identified PCBs included congeners from tri- to decachlorinated, mainly comprising tetra- to octachlorinated PCBs. Of the 59 PCBs, 58 showed connections with all four genes, while the levels of PCB165 were only correlated to *Gstp1*. There were negative correlations between the PCBs and genes *Dpysl2*, *Morf4l1*, and *Spock1*, while the correlations with *Gstp1* were positive. These findings suggest possible mechanistic links between specific PCB congeners detected in the mouse brain and the four DEGs identified in the spatial transcriptomic analysis, which require further investigation.

### PCB exposure impacts BBB integrity

We have previously shown that PCB exposure disrupts BBB integrity [35]. Our spatial transcriptomics data identified the downregulation of *Spock1*, a critical member of BBB development. The structural integrity and permeability of the BBB are facilitated by a variety of proteins, including tight junction proteins and cellular adhesion molecules, which serve to ensure the barrier integrity of this dynamic interface [40, 41]. Occludin, Claudin-5, Zonula Occludens 1 (ZO-1), and Zonula Occludens 2 (ZO-2) are tight junction proteins that restrict paracellular flow while maintaining cell polarity and overall BBB structure [42]. In addition, Afadin contributes to both adherens junction and tight junction integrity. Expression of cell adhesion molecules, such as Intercellular Adhesion Molecule 1 (ICAM-1) and Vascular Cell Adhesion Molecule 1 (VCAM-1), also influences BBB permeability due to their role in regulating the transmigration of immune cells [40]. In this study, qPCR and immunoblotting were performed on whole-brain homogenates of PCB-exposed mice, with vehicle-treated mice as controls. Exposure to the PCB mixture resulted in a significant decrease in protein expression of Occludin and Afadin (**Fig. 7**). Although gene expression of these proteins was not altered, alterations in protein expression are considered a better indicator of long-term changes. In contrast, the expression of Claudin-5, ZO-1, and ZO-2 at either the gene or protein level yielded no notable changes, indicating the specificity of the responses (**Fig. 7**). Although ICAM-1 protein levels exhibited a decreasing trend following PCB exposure, this change was not statistically significant. Similarly, PCB exposure had no discernible effect on VCAM-1 expression at either the gene or protein level (**Supplemental Fig. S4**).

**Figure 7.**
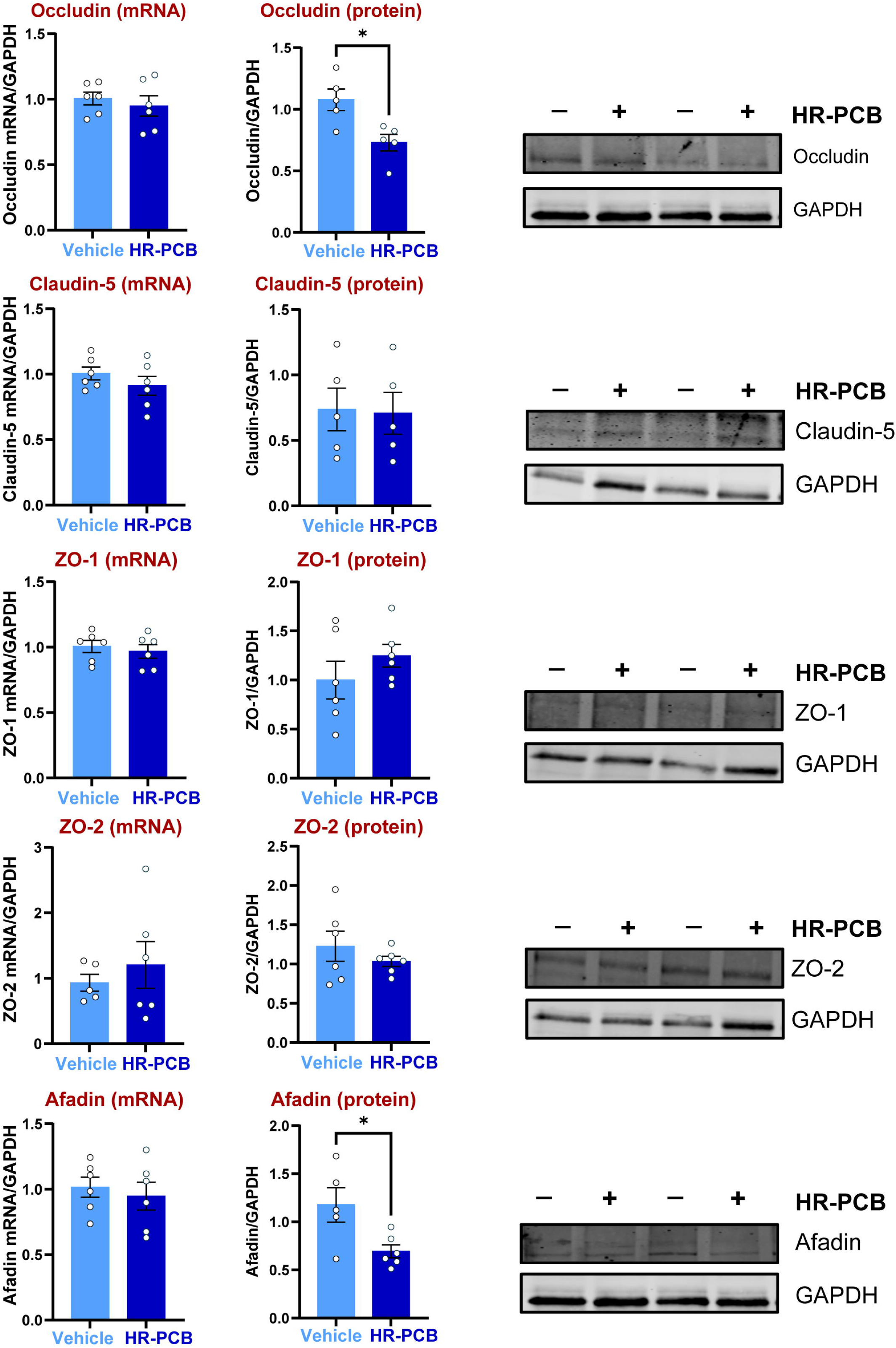
Expression of markers of the BBB integrity. Mice were exposed to the PCB mixture or vehicle control as in Figure 1. Left panels show qPCR data. Middle and right panels show immunoblotting results presented as quantitative bar graphs and representative immunoblots of the target proteins. GAPDH levels were used to normalize the results. Values are mean ±SEM with n = 5-6 per group; *p<0.05.

## Discussion

Cognitive loss, especially spatial memory impairment, is a common symptom of dementia. Epidemiological studies suggest the potential role of environmental toxicants in increasing the risk of ADRD [43]. Particularly, PCB mixtures containing higher chlorinated PCB congeners (>4 chlorine substituents) have been detected in post-mortem human brain samples from older donors [9]. In this study, we demonstrate the impact of exposure to a HR-PCB mixture on spatial memory and transcriptional signature across brain regions in adult mice. We show that daily oral administration of PCB for seven weeks impairs hippocampus-dependent long-term spatial memory. Using a cutting-edge spatial transcriptomics approach, we further delineate the transcriptomic signature impacted by PCB in five brain regions: hippocampus, neocortex, thalamus, caudoputamen, and fiber tracts.

Studies examining the association between serum PCB levels and memory performance in older adults suggest adverse cognitive effects [44]. Studies in monkeys [45] and rats [46] demonstrated that PCB exposure impairs spatial memory. Nevertheless, our findings indicate that the administered dose of HR-PCB does not elicit anxiety-related behaviors nor influence the overall exploration of objects. Therefore, our work demonstrating the impact of HR-PCB exposure on spatial memory impairment in adult mice supports the current literature and provides new insight into the neurotoxic effects of PCBs on spatial memory performance.

Spatial memory closely relies on the hippocampus and its connectivity with the cortex [11]. We and others have shown that gene expression changes in the hippocampus during critical time points following a learning event are essential for spatial memory consolidation [13, 14, 47]. Diseases associated with memory loss, such as ADRD, show pathological tau and amyloid beta deposition in the hippocampus and neocortex [48, 49]. PCBs have been detected across different brain regions, including the hippocampus of post-mortem human brains [9]. Therefore, the spatial memory impairment in PCB-exposed mice could be due to the direct impact of PCBs on the hippocampus and neocortex.

Understanding the impact of PCBs across brain regions requires molecular analyses at high spatial resolution. Advancements in spatial transcriptomic approaches enable the examination of transcriptional signatures across multiple brain regions [14, 18]. Therefore, we used a recently developed spatial transcriptomics approach to investigate the molecular signature of PCB-mediated neurotoxicity in the mouse brain. Our spatial transcriptomic analysis of male mice brains after seven weeks of HR-PCB exposure revealed complex and region-specific, yet overlapping, gene targets across five brain areas: the hippocampus, neocortex, thalamus, caudoputamen, and fiber tracts.

Many genes throughout the brain were impacted by exposure to the HR-PCB mixture. For instance, one gene that was downregulated in the hippocampus and neocortex was *Dpysl2.* This gene is essential for hippocampus-dependent spatial memory [33]. Additionally, we found downregulation of *Tcf4* exclusively in the hippocampus. *Tcf4* is essential for synaptic plasticity and memory consolidation [30], thus providing a possible link between PCB exposure and long-term memory deficits. Interestingly, the upregulation of *Gstp1* in the hippocampus, neocortex, thalamus, and fiber tracts suggests the vulnerability of these brain regions to neurotoxicity caused by PCBs. *Gstp1* is associated with detoxification [38]; thus, increased *Gstp1* expression suggests a neurotoxic impact of PCB in these regions. However, future research on detecting PCB levels at high spatial resolution is essential for understanding the spread of PCB across vulnerable brain regions and cell types.

Among the downregulated genes in the hippocampus and neocortex, *Spock1* plays a crucial role in maintaining BBB permeability. The decrease in the expression of Afadin and Occludin, proteins associated with tight junctions, is in line with these results and suggests BBB dysregulation. While the current analyses were performed on whole-brain homogenates, the results align with previous observations on the microvascular impact of PCB congeners [35]. In addition, they further support conclusions from studies that demonstrated decreased Occludin levels in the hippocampal regions of rats following intraperitoneal injections of PCBs, as well as a study that observed reduced levels of Afadin in human brain endothelial cells treated with PCBs [50, 51]. The adverse functionality of the intricate barrier system, comprised of tight junctions and their cytosolic accessory proteins that regulate the integrity of the brain endothelium in concert, may result in aberrant paracellular movement of ions and solutes across the BBB [52]. This potential disruption is implicated in a range of pathologies, including ADRDs [53]. Indeed, repairing the BBB integrity was demonstrated to have a beneficial impact on the outcomes of depression [54] and stroke [55].

Research on the relationship between specific PCB congeners and gene expression profiles in the brain is limited. A study involving rat liver cells found that exposure to PCB126, a dioxin-like PCB congener [26], significantly increases *Gstp1* expression [56]. PCB126 was not detected in the mouse brains from this study because it is a minor PCB congener present in the HR-PCB mixture. However, other dioxin-like mono-ortho PCBs (e.g., PCB105, PCB118, PCB156, PCB157, PCB167, PCB169, and PCB189) showed positive correlations with *Gstp1* in the hippocampus. These findings suggest an aryl hydrocarbon (AhR) receptor-mediated effect of PCBs on gene expression in the mouse brain, similar to the prototypical effects of dioxin-like compounds on the expression of genes, such as *Gstp1*, in the mouse liver [57]. However, current mechanistic literature does not support a direct AhR-driven neurotoxic pathway for these PCBs [58]. Instead, alterations in calcium signaling, dopamine homeostasis, thyroid hormone interference, or oxidative stress are more likely to explain the neurotoxicity of dioxin-like PCBs. In line with this current mechanistic understanding, the results from the network analyses suggest that the effects of exposure to the HR-PCB mixture on spatial gene expression is a result of the non-AhR effects from higher-chlorinated PCB congeners accumulating in the mouse brain, rather than direct effects mediated by the AhR. However, further mechanistic studies are necessary to validate this hypothesis.

In conclusion, using congener-specific PCB measurements as well as behavioral, biochemical, and state-of-the-art gene expression approaches, we demonstrate the impact of a human-relevant PCB mixture on spatial memory and gene expression changes in the brains of male mice. Our spatial gene expression profiling approach identified the transcriptional impact of HR-PCB across five different brain regions, with persistent, higher chlorinated PCB congeners being associated with changes in spatial gene expression. However, our spatial gene expression data lacks cell-type-specific information. Another limitation of this study is that Visium analysis could cause gene expression bias due to variations in barcodes across tissue regions within the Visium capture area, potentially leading to batch effects in the samples. Multiplexed *in situ* hybridization-based approaches have recently been utilized to determine spatial transcriptional changes at single-cell resolution. Thus, future work should investigate cell-type-specific spatial gene expression analysis following PCB exposure in the brain of male and female mice. These future directions will yield novel mechanistic insights into the sex-specific and cell-type-specific effects of PCB exposure on brain gene expression and function. Such insights will help establish a foundation for developing targeted strategies to prevent or mitigate the neurotoxic effects of PCBs.

## Materials and Methods

### Data reporting

No statistical methods were used to predetermine the sample size.

### Chemicals

Aroclor 1248 (Lot 106-248), Aroclor 1260 (Lot 021-020-1A-01), and 2,3,3’,4,5,6-hexachlorobiphenyl (PCB 160, Lot 29009) were purchased from AccuStandard Inc. (New Haven, CT, USA). Aroclor 1242 (Lot KB-05-415), and Arolcor 1254 (Lot KC-12-638) were obtained from the Synthesis Core of the Iowa Superfund Research Program. 2,4,4′-Trichlorobiphenyl (PCB 28) was synthesized via a Suzuki coupling reaction of 4-chlorobenzene boronic acid with 2,4-dichlorobromobenzene [59].

### HR-PCB mixture preparation

A human-relevant PCB (HR-PCB) mixture that simulates the average human brain PCB profile measured in the cerebellum from 30 male and 42 female donors, aged 8 to 59 years (average age, 33 years) (**Supplemental Fig. 1A**). Additional details regarding the preparation and authentication of the synthetic HR-PCB mixture are available on the Iowa Research Online platform [60]. Briefly, the PCB mixture was prepared from commercial Aroclor mixtures and two individual PCB congeners. First, a theoretical mixture that approximates the target PCB profile was simulated using published profiles of different Aroclor PCB mixtures [61] with the addition of individual PCB congeners [61] (**Supplemental Fig. 1B**). Second, based on this simulation, the synthetic HR-PCB mixture was prepared with the following composition, expressed as a mass percentage: Aroclor 1242 (5%), Aroclor 1248 (19%), Aroclor 1254 (30%), Aroclor 1260 (32%), PCB 28 (8%), and PCB 160 (6%). The similarity between the PCB profiles was characterized using the similarity coefficient cos θ, which ranges from 0 to 1, where a value of 0.0 describes two completely different profiles, and 1.0 describes two identical profiles. The PCB profiles of the theoretical (**Fig. 2B**) and the synthetic HR-PCB mixtures (**Fig. 2C**) were similar to the average human brain PCB profiles, with cos θ values of 0.92 and 0.91, respectively.

### Animals

#### ARRIVE guidelines were followed in reporting materials and methods used throughout the animal study

Twenty-four adult male C57BL/6J male mice were purchased from Jackson Laboratories (#000664) and, at 35 days of age, exposure to PCB or vehicle was initiated. All animals were housed under a standard 12-hour light/12-hour dark cycle with free access to food and water. All the experiments were conducted in accordance with U.S. National Institutes of Health guidelines for animal care and use. They were approved by the Institutional Animal Care and Use Committee of the University of Iowa, Iowa. The animal was considered the statistical unit. No adverse effects were observed throughout the study. However, two animals (one PCB and one vehicle) were removed from behavioral analysis due to being statistical outliers (greater than two standard deviations from the mean).

### PCB preparation and dosing

Male mice were randomly assigned to pairs and placed in cages with corn cob bedding. Compared to common paper bedding, corn cob bedding (Teklad 7907 Irradiated ¼” Corn Cob Bedding) allowed for easier introduction of peanut butter exposures. One plastic hut (product numbers K3102 or K3272, Bio-Serv, Flemington, NJ, USA) was placed in each cage as enrichment. After arriving at the University of Iowa animal housing facility, mice were given one week to acclimate. In the second week, mice were acclimated to a peanut butter (Trader Joe’s)/peanut oil (Spectrum Organic Products) mixture without PCBs to train them to consume the peanut butter. Exposures were performed at Zeitgeber time (ZT) 6 (±1 hour) to maintain a consistent behavioral pattern and avoid circadian biases [62]. Weights in grams were collected for each mouse the day before feeding and used to calculate the dose of peanut butter or peanut oil administered (0.05 g PB/PO per 40 g body weight). The samples were placed in a small weigh boat for easy delivery. At the time of feeding, the huts were removed, and a custom polycarbonate divider was carefully inserted into the mouse cages to split the cage area in half, with one mouse in each half. Then, the appropriate sample was given to each mouse. The mice were observed to ensure that the sample was eaten, and the time taken to consume it was recorded. Dividers were removed and cleaned when feeding was complete, and the huts were reintroduced. At the start of the third week, mice were given either 6 mg/Kg body weight PCB mixture in a peanut butter/peanut oil mixture or peanut butter/peanut oil alone for seven weeks. Exposures were performed daily until SOR training was performed.

### PCB Extraction from the Brain Tissue

PCBs were extracted from the brain using liquid-liquid extraction, adapted from a method as previously described [63]. Briefly, approximately 60 mg of brain tissue (62 ± 11 mg, n=4 mice/treatment group) was homogenized in 3 mL of 2-propanol using a TissueRuptor (QIAGEN, Hilden, Germany). After spiking all samples with stable isotope ^13^C labelled surrogate standards (10 standards to represent each homolog, 10 ng each), PCBs were extracted using a 1:9 (v/v) mixture of diethyl ether and hexane. The extracts were then washed with 5 mL of 0.1 M phosphoric acid in 0.9% sodium chloride solution. Following concentration under a gentle stream of nitrogen, the extracts were passed through a cartridge preloaded with acidified silica gel (sulfuric acid : silica gel, 1 : 2, w/w) to remove the lipid. Finally, the extracts were again concentrated under nitrogen and spiked with internal standards (d-PCB30 and PCB204, 10 ng each) for GC-MS/MS analysis.

### GC-MS/MS Instrumental Setup for PCBs Analysis

PCBs were quantified using a GC-MS/MS system (Agilent 7890B GC, 7000D Triple Quad, and 7693 autosampler; Agilent Technologies, Santa Clara, CA, USA), following methods described previously [64, 65]. The system was equipped with a Supelco SPB-Octyl capillary column (50% n-octyl/50% methyl siloxane, 30 m length, 0.25 mm inner diameter, and 0.25 µm film thickness; Supelco, Bellefonte, PA, USA). Helium served as the carrier gas at 0.8 mL/min, and nitrogen was used as the collision gas. The gas chromatograph operated in solvent vent injection mode (initial temperature of 45 °C for 0.06 min, followed by a rapid ramp of 600 °C/min to reach an inlet temperature of 325 °C at 5 psi). The GC oven temperature program began at 45 °C for 2 min, then increased to 100 °C at a 160 °C/min, then to 250 °C at 1.6 °C/min, and finally increased to 280 °C at 80 °C/min, with a final hold time of 11 min. The transfer line temperature was 280 °C, and the electron ionization source temperature in the triple quadrupole was maintained at 230 °C. The quantification of PCBs was based on the relative peak areas compared to internal standards, with adjustments for surrogate recoveries. The PCB precursor and product masses of unlabeled and ^13^C-labeled calibration standards employed in multiple reaction monitoring mode are reported in **Supplemental Table S1**. PCB levels were adjusted for tissue wet weight. In addition, the 2,3,7,8-tetrachlorodibenzodioxin (TCDD) toxic equivalents were calculated using this formula: TEQ=∑(PCB_i_ ×TEF_i_), where PCB_i_ indicates the levels of PCB (in ng/g) and TEF_i_ is the TEF value for the corresponding PCB [26].

### Quality Assurance and Quality Control (QA/QC) for PCB Analyses

Solvent blanks, method blanks, and laboratory reference material (LRM) were processed alongside all samples to evaluate the reproducibility and precision of the analyses. The LRM was prepared with pooled and homogenized brain samples from a previous study that exposed female rats to a PCB mixture via nose-only inhalation system [66]. The measured levels of PCBs were adjusted based on the recoveries of their respective surrogate standards. The method detection limit (MDL) was determined using the formula: MDL = mean_blank_ + t_(0.01, n-1)_ × SD_blank_, where mean_blank_ is the average concentration in the method blanks, t_(0.01, n-1)_ is the Student’s t-value for n–1 degrees of freedom at a 99% confidence level, and SD_blank_ is the standard deviation of the method blanks [67]. Similarly, the limit of detection (LOD) was calculated using the formula: LOD = mean_control_ + t_(0.01, n-1)_ × SD_control_, where mean_control_ and SD_control_ represents the average concentration and standard deviation, respectively, obtained from control tissue samples. The surrogate standard recoveries, MDLs, LODs, and LRM (including average, standard deviation and relative standard deviation) are reported in **Supplemental Tables S2-S5**.

### Spatial object recognition task

The SOR task was used to assess long-term memory at approximately 3.5 months of age in male C57BL/6J mice. After the final day of PCB exposure, all mice were exposed to an open field for 6 minutes during habituation, without any objects. (HR-PCB, n=11, and vehicle, n=11). The sample size was determined based on previous publications that assess long-term spatial memory using SOR task [14]. After habituation, mice were returned to their home cage, and the open-field chambers were cleaned. Two clean glass objects were placed in a specific spatial location in the open field, and mice were introduced back with the objects. Training with objects was continued for 6 minutes, and then the mice were returned to the home cage while objects and the open fields were cleaned with 70% ethanol. Three training sessions with objects were performed. Long-term memory was assessed after 24 hours by moving one object to a novel spatial location. Exploration time of the objects was recorded for each session and calculated to show discrimination towards the displaced object using the calculation:

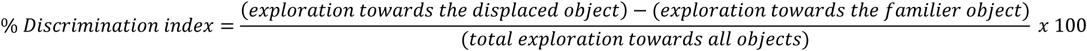

### Sample preparation for spatial transcriptomics

Mouse brains were collected one hour after the testing in the SOR task. The mice were euthanized by cervical dislocation, and the whole brains were flash-frozen using −70^0^C isopentane. Flash-frozen brains were stored at −80C until sectioning. Brains were adhered in optimal cutting temperature medium (OCT) and then cryosectioned at −20°C with the Leica CM3050 S Cryostat. Coronal sections measuring 10 microns thick were prepared from the brain area containing the dorsal hippocampus. They were placed on chilled Visium Spatial Gene Expression Slides (10X Genomics). Trimmed coronal sections of two biological replicates from the same group were mounted in each capture area of the Visium slide. The trimming of the coronal sections was performed in such a way that the sections remained centered around the hippocampal region. The Visium slides were fixed, stained, and imaged after being stained with Hematoxylin and Eosin (H&E) on an Olympus BX61 microscope. The tissue was subsequently permeabilized, then Poly-A mRNAs captured by the poly(dT) primers pre-coated on the slide. Reverse transcription and synthesis of the second strand were employed to amplify cDNA samples from the Visium slides. These samples were subsequently transferred, purified, and quantified for library preparation. Sequencing libraries were prepared according to the Visium Spatial Gene Expression User Guide (10X Genomics) by the Iowa Institute of Human Genetics (IIHG) Genomics Division. The libraries were sequenced on a NovaSeq 6000 (Illumina).

### Spatial transcriptomics analysis

Space Ranger 3.0.0 was used for mapping the raw fastq files to the mouse reference genome and aligning the sequencing data to the microscope image of H&E-stained coronal sections. Following sequence alignment with the microscopic image, the replicate tissue sections were selected using the lasso tool in the loupe browser. The spatial coordinates for the selected cells were exported from the loupe browser and were used to subset the replicates from the composite Seurat object. Four replicates of vehicle and PCB treated samples were individually saved as separate Seurat objects. SCTransform normalization was performed on each replicate separately to account for library size differences across the Visium spots. The replicates from vehicle and PCB treated samples were integrated using the Seurat integration pipeline. Briefly, the integration anchors were chosen (FindIntegrationAnchors) from the list of eight Seurat objects of vehicle and PCB exposed samples. These anchors were then used to integrate the eight datasets together (IntegrateData). Linear dimensionality reduction was performed on the integrated Seurat object by principal component analysis (runPCA, npcs = 30). A k-nearest-neighbors graph was constructed based on Euclidean distance in PCA space and refined (FindNeighbors), following which the Visium spots were clustered using Louvain algorithm at resolution 0.1. Clusters were visualized with UMAP. Brain regions were annotated based on their spatial location and marker gene expression based on Allen Brain Atlas [68]. Fisher’s exact test was performed to evaluate the change in proportion of barcoded spots (replicates pooled together) in each brain region between vehicle and PCB exposed groups (**Supplemental Table S6**).

### Differential gene expression

Differential gene expression analyses per brain region between vehicle and PCB exposed groups were performed on the scaled expression data using findMarkers (assay = ‘SCT’, test.use = ‘wilcox’). The genes with a false-discovery rate (FDR) < 0.05 and |log2 fold-change| > 0.2 were considered significant (**Supplemental Table S7**).

### Gene ontology enrichment analysis

The DEGs were subsequently sorted to find out the genes that were uniquely upregulated or downregulated in each brain region. The common genes that were differentially expressed in more than one brain regions were also extracted. Those unique and common genes were subjected to Molecular Function (GO:MF) enrichment analysis using the ‘enrichGO’ function in the clusterProfiler package with the following criteria: pvalueCutoff = 0.01 and qvalueCutoff = 0.05. The GO:MF enrichment analysis was performed for the upregulated and downregulated gene sets separately (**Supplemental Table S8-9**). The unique and common DEGs for each brain region were visualized using ComplexHeatmap and the top molecular functions enriched for the list of genes were annotated in the heatmap.

### Multi-omics Network Analysis

A comprehensive network analysis was conducted to investigate the interactions between PCB congener levels and DEGs across five brain regions. The pseudo-bulk gene expression profile of DEGs (shown in Figure 4) across the five brain regions was extracted using the Seurat AggregateExpression(assays = "SCT") function. Network analysis was performed with xMWAS (version 1.0, https://kuppal.shinyapps.io/xmwas/), utilizing the partial least squares (PLS) regression analysis and eigenvector centrality measures [69]. The thresholds of absolute correlation coefficients R were set at > 0.7 for neocortex and > 0.9 for hippocampus, with significance determined by Student’s t-test (p < 0.05). To aid interpretation, a sub-network connecting four selected DEGs, including *Spock1*, *Dpysl2*, *Morf4l1*, and *Gstp1*, was extracted and highlighted (**Fig. 6**). These four genes were not identified in the thalamus, caudoputamen, or fiber tracts even with a less stringent threshold of R > 0.4. Final network visualization and annotation were carried out using Cytoscape (version 3.10.1) [70].

### RT-qPCR

Total RNA was extracted from whole brain homogenates using the RNeasy Lipid Tissue Kit (Qiagen) followed by reverse transcription using the qScript XLT 1-Step RT-qPCR ToughMix Low ROX (Quantabio). Nucleic acid concentrations were determined using a Nanodrop 2000. A total of 100 ng of RNA was used for the reactions, with GAPDH (VIC; Thermo Fisher) as an endogenous control. The following primers were utilized with the QuantStudio 6 Flex Real-Time PCR System (Thermofisher Scientific): Mm00500910_m1 (occludin); Mm00727012_s1 (claudin-5) Mm00516023_m1 (ICAM1); Mm01320970_m1 (VCAM1); Mm01320638_m1 (ZO-1), Mm00495620_m1 (ZO-2), Mm01273324_m1 (afadin).

### Western blotting

Whole brain tissue was homogenized using rapid immunoprecipitation (RIPA) lysis buffer (Fisher Scientific) plus protease and phosphatase inhibitor cocktail (Fisher Scientific), followed by centrifugation, and protein quantification using the Pierce BCA assay kit (Fisher Scientific). Samples were diluted in RIPA and sodium dodecyl sulfate (SDS) and 50 µg per sample was loaded into a 4-20% midi-PROTEAN TGX Stain-Free polyacrylamide precast gel (Bio-Rab Laboratories). Samples were then transferred to a nitrocellulose membrane using the Trans-Blot Turbo System (Bio-Rad Laboratories). Membranes were blocked overnight at 4°C using 5% bovine serum albumin (BSA) and then incubated with primary antibodies diluted in 5% BSA overnight at 4^0^C. Blots were washed three times using Tris-buffered saline with 0.1% Tween 20 (TBS-T) and incubated with secondary antibody at room temperature for 1 h as follows: IRDye 680RD Goat anti-rabbit (1:10,000; Licor) and IRDye 800CW Goat anti-mouse (1:10,000; Licor). GAPDH (1:10,000; Invitrogen) was utilized as an endogenous control and for sample normalization. Bands were imaged using the Odyssey CLX Imaging System (Licor) and the signal was quantified using the Image Studio 4.0 software (Licor).

### Statistical analysis

The text indicates statistical analyses, using the animal as the statistical unit, and graphs were prepared using GraphPad Prism (Version 10.5.0). Statistical outlier limits were determined as greater than two standard deviations from the mean. Statistical analyses were performed using either a 2-tailed Student’s t-test or a 2-way ANOVA followed by Sidak’s multiple comparisons post hoc tests. The factors considered for 2-way ANOVA were session (training or test) and exposure (PCB or vehicle). The mean of all three training sessions was considered as training. In all cases, differences were considered significant when p ≤ 0.05. Error bars in all figures represent ± standard error (SEM).

## Acknowledgments

We thank the Neural Circuits and Behavior Core, Iowa Neurobank Core, and the Iowa Institute of Human Genetics (IIHG) Core for using their facilities, Julia Neuharth for technical assistance, Dr. Keri C. Hornbuckle for support from the Iowa Superfund Research Program. Human tissue used to develop the HR-PCB mixture was received from the NIH NeuroBioBank at the University of Miami and the Sepulveda Research Corporation.

## Funding

This work was supported by grants from the National Institute of Health R01 ES014901, R01 ES031098, and R01 ES034691 to H-J.L., and ES036983 to M.T. D.T was supported by a grant from the Howard Hughes Medical Institute through the Gilliam Fellows Program. The synthesis of the HR-PCB mixture and the PCB analyses was performed in facilities of the Iowa Superfund Research Program (P42 ES013661) and the Environmental Health Sciences Research Program (P30 ES005605) at the University of Iowa The sequencing data presented herein were obtained at the Genomics Division of the Iowa Institute of Human Genetics (RRID: SCR_023422), which is supported, in part, by the University of Iowa Carver College of Medicine.

## Author contributions

S.C. and H-J.L. conceived the idea and supervised the work. S.C., H-J.L. and B.B. wrote the manuscript with input from all the authors. S.E.B., S.L., and Z.N. dosed PCB, performed the behavior, and analyzed behavioral data. B.B. performed the analysis of the spatial transcriptomics data. N.M.B. and L.D. prepared the PCB dosing mixtures. X.L. prepared and authenticated the HR-PCB mixture. H.W. performed the brain PCB level assessment and transcriptomics network analysis. R.F.M. supported the PCB measurements. M.J.T. and D.T. performed biochemical analysis and interpreted the data.

## Competing interests

The authors declare no conflicting interests.

## Supplementary information

Supplementary Information includes Supplemental Figures, Supplemental Figure legends 1-4, and Supplemental Tables legends S1-9.

## Data availability

The spatial transcriptomic datasets produced in this study are available in the NCBI Gene Expression Omnibus (GEO) database under the accession code GSE301012.

## Code availability

The custom scripts used for analyses and generating figures related to spatial transcriptomics data can be accessed through GitHub (https://github.com/ChatterjeeEpigenetics/PCB_Brain_Visium.git).

## Notes

### Competing Interest Statement

The authors have declared no competing interest.

