## Supplemental information for "Spatial transcriptomic profiling uncovers the molecular effects of the neurotoxicant polychlorinated biphenyls (PCBs) in the brains of adult mice"

### Supplementary Information

Supplemental Figures, Supplemental Figure legends S1-4, and Supplemental Tables legends S1-9.

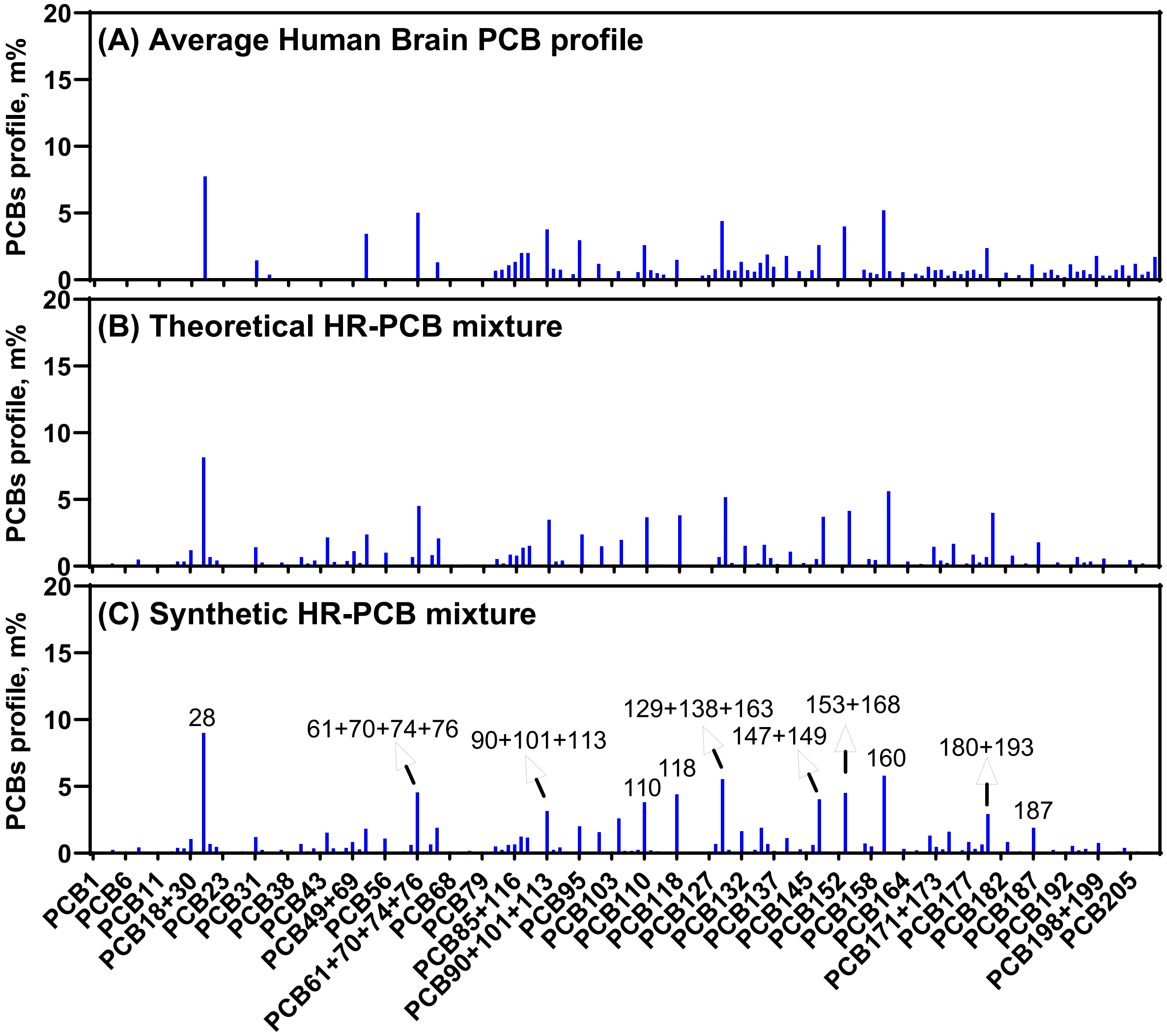


**Supplemental Figure S1.** Comparison of PCB profiles (**A**) detected in postmortem human brain tissue with (**B**) the theoretical HR-PCB mixture developed based on mixing technical Arcolor PCB mixtures and individual PCB congeners, and (**C**) the synthetic HR-PCB mixture prepared to match the theoretic PCB profile. The human brain PCB profile represents the average PCB profile in the cerebellum from 30 male and 42 female donors, aged 8 to 59 years (average age, 33 years). The PCB profile shown in panels (A) and (C) were determined by GC-MS/MS. Human tissue was obtained from the NIH Neurobiobank at the University of Maryland, Baltimore, MD. See the details of the data in dataset (Li, X. et al. 2025 https://doi.org/10.25820/data.007553).

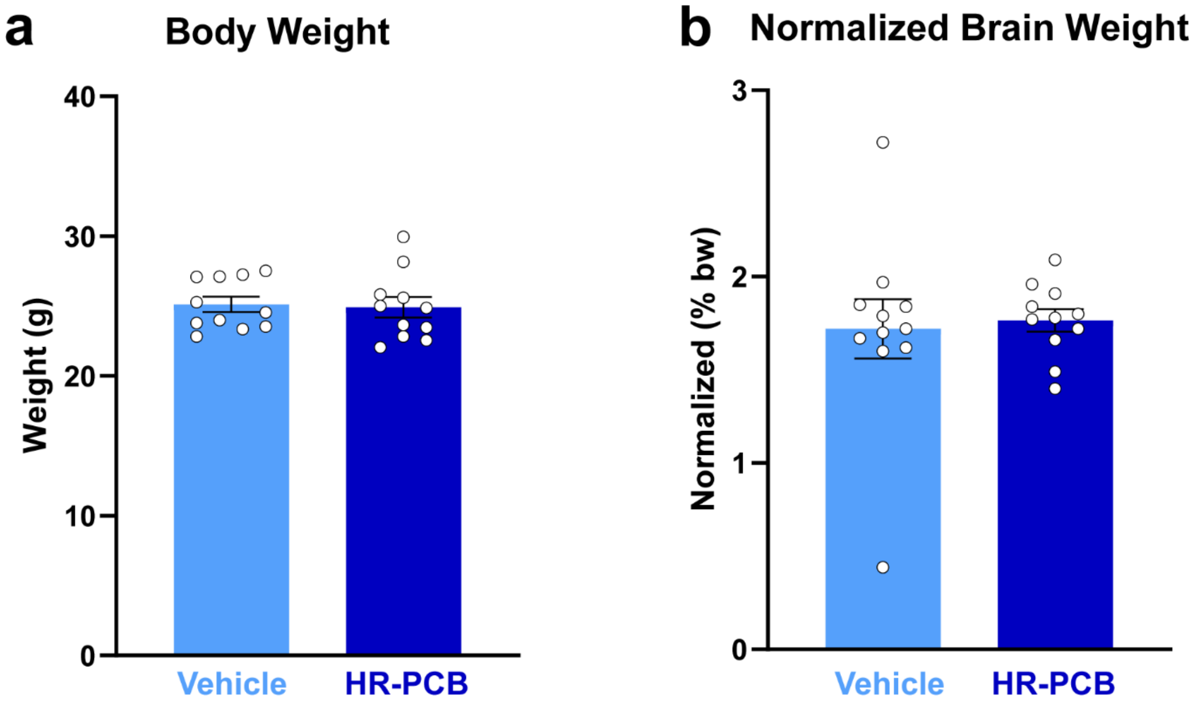


**Supplemental Figure S2. Body and brain weight assessment in PCB and control animals.** Body weight taken from last day of PCB exposure. Wet weight of brain adjusted for bodyweight of mouse. Error bars represent ± SEM. Vehicle (n=11), and HR-PCB (n=11).

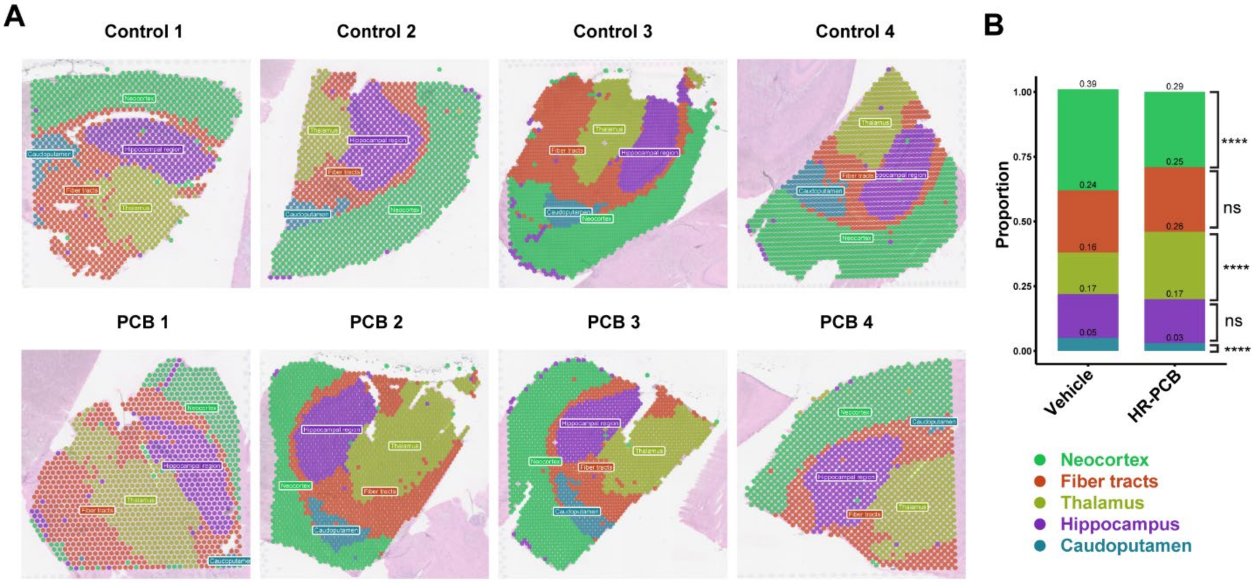


**Supplemental Figure S3. Brain section replicates for the Visium experiment. A.** Barcode spots (colored according to brain regions) overlayed on H&E-stained brain sections are shown for each biological replicate within each group. **B.** Bar plots showing the proportion of visium spots across different brain regions between PCB and vehicle-exposed samples. Fisher exact test comparing vehicle vs PCB: Neocortex ****p-adj=1.78E-30, Fiber tracts p-adj=0.159 (ns), Thalamus ****p-adj=1.53E-41, Hippocampal region p-adj=0.645 (ns), and Caudoputamen ****p-adj=2.67E-05. Vehicle (n=4 mice), and HR-PCB (n=4 mice).

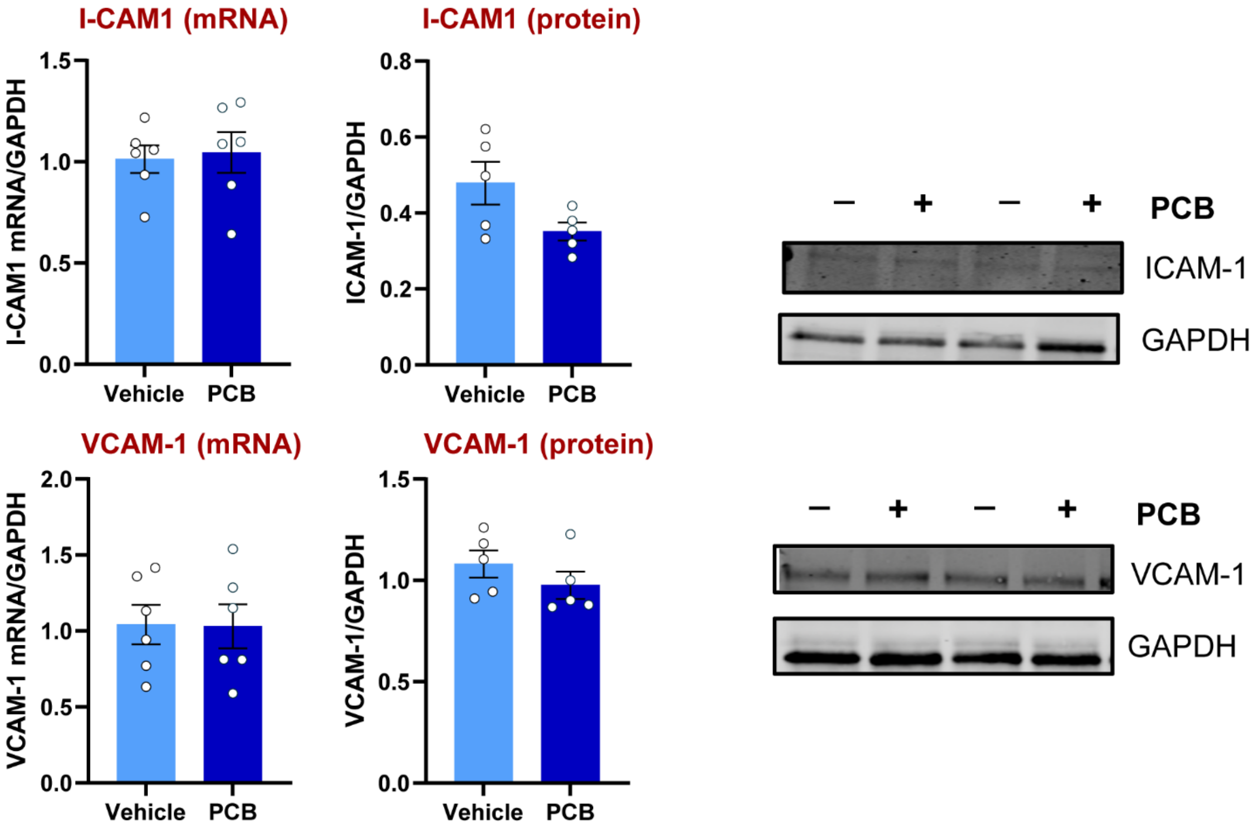


**Supplemental Figure S4. Expression of adhesion molecules ICAM-1 and VCAM-1.** Analyses were performed in whole brain homogenates of mice exposed to the PCB mixture or vehicle control as in Figure 1. Left panels, RT-qPCR data. Middle and right panels, immunoblotting results presented as quantitative bar graphs and representative immunoblots of the target proteins. GAPDH levels were used to normalize the results. Values are mean ±SEM with n = 5-6 per group.

**Supplementary Table legends**

**Supplemental Table S1.** PCB precursor and product masses of unlabeled and $^{13}C$-labeled calibration standards employed in multiple reaction monitoring mode on the triple quadrupole mass spectrometer. Unlabeled standards were from AccuStandard, New Haven, CT, USA. Labeled standards were from Cambridge Isotope Laboratories, Inc.

**Supplemental Table S2.** Surrogate standards recoveries for $^{13}C$ labeled PCBs for each homolog group.

**Supplemental Table S3.** Method Detection Limit (MDL) for each PCB congener or co-eluted congeners. MDLs were calculated using method blanks and expressed as the upper limit of 99% confidence interval (average + $t_{n-1}$* standard deviation, $t_{n-1}$ represents Student's t-value of 99% confidence level with n-1 degree of freedom). Values are expressed in ng.

**Supplemental Table S4.** Limit of Detection (LOD) for each PCB congener or co-eluted congeners. LODs were calculated using control/blank tissues and expressed as the upper limit of 99% confidence interval (average + $t_{n-1}$* standard deviation, $t_{n-1}$ represents Student's t-value of 99% confidence level with n-1 degree of freedom). Values are expressed in ng/g tissue.

**Supplemental Table S5**. PCB levels in the laboratory reference material (LRM). N=7. RSD=SD/mean*100.

**Supplemental Table S6.** Change in proportion of barcoded spots in each brain region between vehicle and PCB exposed groups (Fisher exact test).

**Supplemental Table S7.** Differentially expressed genes in five brain regions (neocortex, hippocampus, thalamus, fiber tracts, and caudoputamen)

**Supplemental Table S8.** Gene Ontology (Molecular Function) enrichment terms on shared and uniquely upregulated genes in each brain region.

**Supplemental Table S9.** Gene Ontology (Molecular Function) enrichment terms on shared and uniquely downregulated genes in each brain region.
